# Chemoproteomics Enabled Discovery of Selective Probes for NuA4 Factor BRD8

**DOI:** 10.1101/2021.02.22.432362

**Authors:** David Remillard, Nikolas A. Savage, Alexia T. Kedves, Joshiawa Paulk, Xin Chen, Francisco J. Garcia, Michael J. Romanowski, Patricia A. Horton, Jason Murphy, Markus Schirle, Edmund M. Harrington, Matthew B. Maxwell, Helen Trinh Pham, Igor Maksimovic, Jason R. Thomas, William C. Forrester

## Abstract

Bromodomain-containing proteins frequently reside in multisubunit chromatin complexes with tissue or cell state-specific compositions. Recent studies have revealed tumor-specific dependencies on the BAF complex bromodomain subunit BRD9 that are a result of recurrent mutations afflicting the structure and composition of associated complex members. To enable the study of ligand engaged complex assemblies, we established a chemoproteomics approach using a functionalized derivative of the BRD9 ligand BI-9564 as an affinity matrix. Unexpectedly, in addition to known interactions with BRD9 and associated BAF complex proteins, we identify a previously unreported interaction with members of the NuA4 complex through the bromodomain-containing subunit BRD8. We apply this finding, alongside homology model guided design, to develop chemical biology approaches for the study of BRD8 inhibition, and to arrive at first-in-class selective and cellularly active probes for BRD8. These tools will empower further pharmacological studies of BRD9 and BRD8 within respective BAF and NuA4 complexes.

## Introduction

The bromodomain is an evolutionary conserved motif that plays important roles in chromatin organization, remodeling and gene control.^1-2^ This domain is the prototypical reader that binds selectively to histones and chromatin regulators that have been post-translationally modified by lysine acetylation (KAc). Its role as a reader serves to establish protein-protein interactions that are fundamental to transcriptional regulation and associated cell states. In this role, the human complement of 46 bromodomain-containing proteins are integral to many multi-subunit chromatin complexes that perform key regulatory transactions within the context of nucleosomal DNA. The discovery of JQ1 as a small molecule probe of BRD4 and the BET family inspired detailed studies of bromodomain proteins and has spurred small molecule inhibitor development throughout the domain family.^3^

The Bromodomain’s antiparallel a-helix bundle anchors KAc residues to an H-bond donating asparagine residue within a central hydrophobic cavity.^4^ This pocket offers an amenable site for KAc competitive inhibition or targeted protein degradation approaches.^3, 5-7^ Such probes offer a point of intervention in the dysregulated gene regulatory networks associated with cellular disease states.^8-9^

Previously, we developed heterobifunctional degradation probes for bromodomain-containing protein 9 (BRD9), a subunit of the BAF (SWI-SNF) chromatin remodeling complex.^6^ These probes enabled validation and study of a remarkable cancer-restricted BRD9 dependency in the context of synovial sarcoma, and have initiated the development of clinical candidates for this disease. ^6, 10-12^ In synovial sarcoma, a precisely fused SS18-SSX oncogenic BAF subunit drives dysregulated chromatin signaling, a process that renders addiction to the otherwise non-essential BRD9 bromodomain subunit. Similarly, BRD9 dependency has also been linked to rhabdoid tumor, another soft tissue cancer that also involves a dysregulated BAF assembly owing to loss of function mutations in the SMARCB1 subunit.^13-14^ These findings have provided a broader insight that the assembly of contextually varied BAF subunits is key to an informed understanding of target biology for BRD9, and motivate the study of such co-complex context effects for both BRD9 and other bromodomain subunits that exist within chromatin complexes.

To enable study of the target profile and the associated non-covalent assembly engaged by BRD9 probes, we establish herein a chemoproteomic affinity enrichment strategy in the native proteome of the BRD9-sensitve acute myeloid leukemia (AML) lineage THP1.^15^ Unexpectedly, we find that in addition to BAF, the widely used BI-9564 BRD9 probe engages the native NuA4 acetyltransferase complex via a previously unreported off-target activity against the known bromodomain-containing BRD8 (p120) subunit.^16^ BRD8 has been previously linked to NuA4’s DNA repair activity,^17-18^ and implicated as a target in colorectal and hepatocellular carcinoma, ^19-21^ as well as a regulator of the responsiveness to DNA damaging chemotherapy.^20^ However, to date no small-molecule inhibitors of BRD8 have been reported. We therefore leverage this finding to develop assays for the study of BRD8 bromodomain inhibition, and apply model-guided design to develop selective first-in-class inhibitors of this understudied NuA4 KAc reader.

## Results

To map the proteome-wide binding profile of BRD9 targeted bromodomain probes in native cell lysates, we applied a chemoproteomics strategy in which protein targets are captured by a probe-functionalized affinity matrix, before their identification and quantification by mass spectrometry. In this approach, pre-treatment of lysates with free probe results in competition of specifically enriched targets, along with tightly associated binding partners, which can be quantified as loss of signal using Tandem Mass Tag (TMT)-multiplexed reporter ion intensities (Figure 1A). To apply this technique to profile targets of BI-9564, we prepared compound **<u>1</u>**, an analog featuring a primary amine to enable direct conjugation to solid support via NHS-ester coupling (Figure 1B). Using the resulting affinity matrix, chemoproteomic experiments were performed in duplicate where TMT tagging provided relative protein abundances from vehicle (DMSO) versus free compound **<u>1</u>** pretreated enrichments from lysate of THP1 cells, a lineage with known dependence on BRD9.^15^

**Fig 1.**
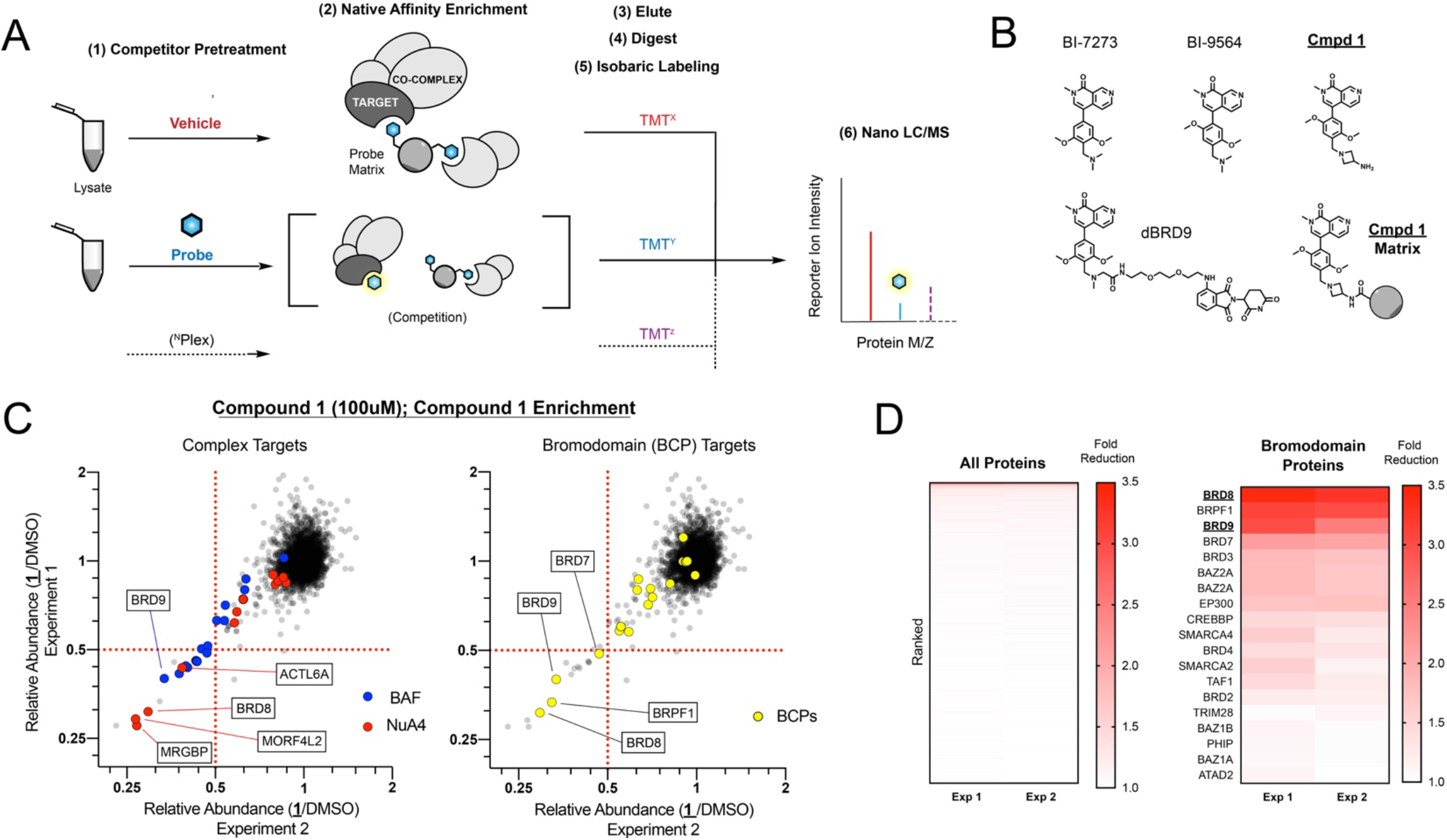
Chemoproteomics Identifies unexpected off target binding to the NuA4 complex. (A) Schematic of the chemoproteomics workflow. (B) Structures of literature compounds and chemoproteomic tool Compound **<u>1</u>**. (C) Scatterplot view of duplicate chemoproteomics with Compound **<u>1</u>** competition (100μM), highlighting NuA4 and BAF complexes (left) or bromodomain-containing protein (BCP) targets (right). (D) Ranked heatmap of fold competition across all identified proteins (left) versus BCP targets (right).

Within the >3000 proteins identified with standard filtering criteria from two replicates, 18 proteins were identified as robustly competed by pretreatment with 100µM free compound **<u>1</u>** (≥50% loss in two replicates), reflecting a generally narrow proteomic binding profile for this probe (Figure 1C,1D). The high potency target BRD9 and its associated BAF co-complex members were identified as selectively competed from the affinity matrix as anticipated. In addition, four members of the NuA4 complex were identified within this competed protein subset, suggesting that an intact assembly of this acetyltransferase complex was engaged by compound **<u>1</u>** (Figure 1C). Collectively, components of the BAF and NUA4 complexes comprised 13 of 18 robustly competed targets, alongside two MOZ/MORF complex members, ING1 and associated bromodomain subunit BRPF, a minor off-target of the parental BRD9 probe.^22^ Among the identified NuA4 members, we noted the dual-bromodomain-containing protein BRD8 as a likely site for recruitment to the affinity matrix due to its potential as an off-target interaction of compound **<u>1</u>**. BRD8 was not included in previous family-wide profiling efforts, and scored as the top competed target among all identified bromodomain-containing proteins in this study (Figure 1D).

To test the possibility that BRD8 is directly engaged by **<u>1</u>**, we prepared a biotin-functionalized “tracer” analog of this compound that would enable the evaluation of direct binding in-vitro with recombinantly expressed BRD8 bromodomain (Figure 2A). For these studies, we initially considered the amino terminal bromodomain of BRD8 (BRD8(1)) isolated as a his-tagged fusion protein. Reconstitution of these components in an AlphaScreen format generated a robust luminescence signal reflecting the extent of tracer-target engagement, and upon further optimization provided a competitive dose response format for use in structure activity relationship (SAR) studies alongside similar assays for BRD9 and BRD4 (Figure 2B).^6, 23^ We observed that binding of the tracer to BRD8 was competed with moderate potency by parental probe BI-9564 (IC50 = 970 nM) and by unmodified compound **<u>1</u>**, less potently by I-BRD9 and BI-7273, and not competed by the BET bromodomain probe JQ1 (Table 1). These data provide supporting biophysical evidence that the first bromodomain of NuA4 complex member BRD8 is an off-target of BI-9564 and other structurally related BRD9 bromodomain probes.

**Table 1.**
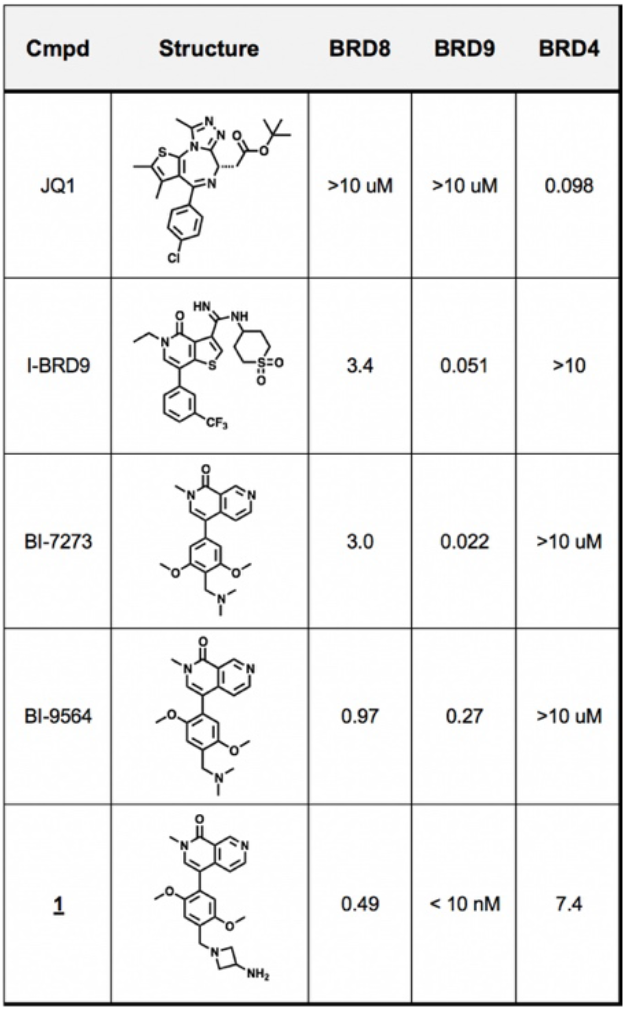
AlphaScreen Characterization of Literature and Chemprot tool <u>1</u>.

**Fig 2.**
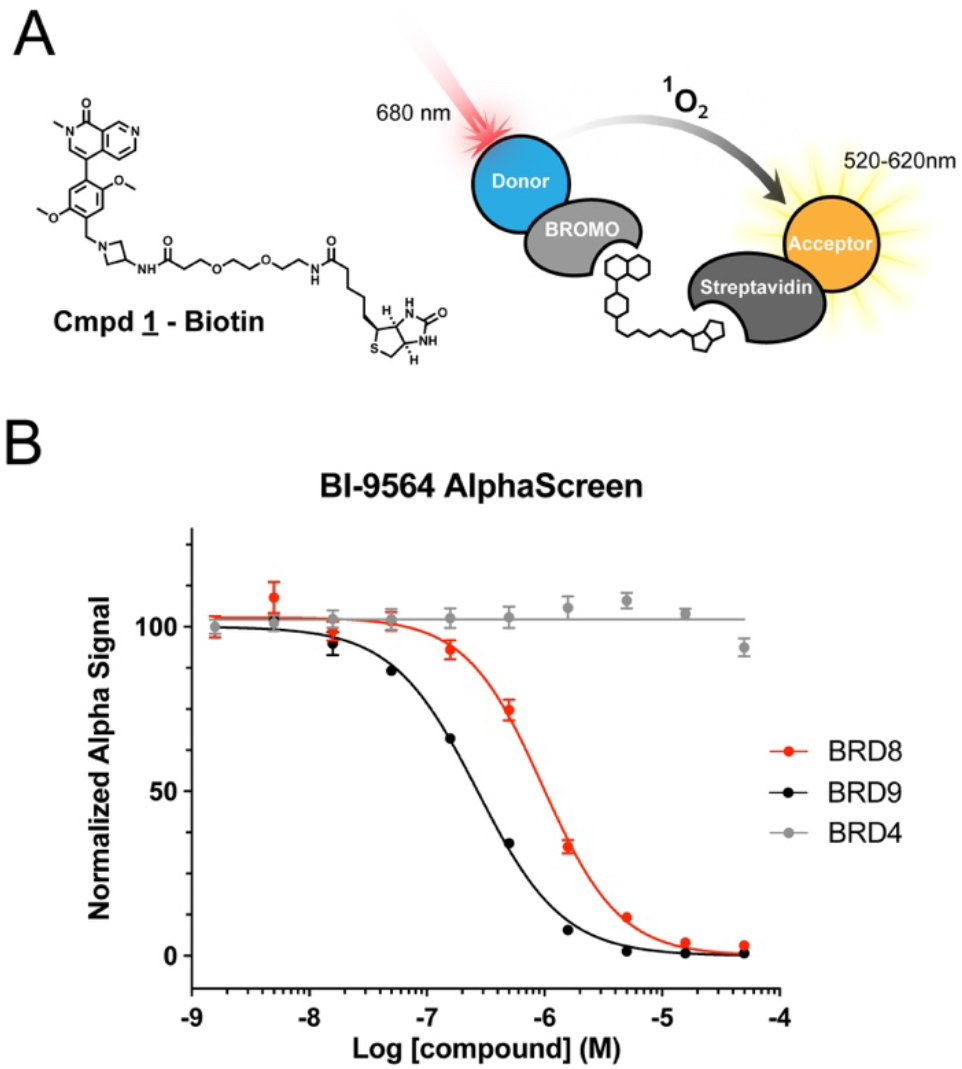
Implementation of a BRD8 AlphaScreen assay to examine bromodomain binding profiles in vitro. (A) Structure of BRD8 AlphaScreen tracer (Compound <u>1</u>-Biotin) (left) and assay schematic (right). (B) Representative AlphaScreen competition curves for literature compound BI-9564.

As no small molecule probes have been previously reported for BRD8, we sought to establish a binding model of BI-9564 and BRD8(1), which might inform opportunities for optimization of potency and selectivity toward a first-in-class BRD8 tool compound. BRD8’s dual bromodomains have proven difficult to express in crystallographic quality and quantity within the efforts of our group, and the attempts of others,^24^ and currently no structural data for these domains is available. In the absence of an existing structure, we sought to generate a model for the binding of BI-9564 to BRD8(1) by homology modeling using a previously solved BI-9564-BRD9 co-crystal structure (PDB ID: 5F1H) as a template.

Given the high degree of structural conservation across the bromodomain family, we were optimistic about the adaptability of this in silico approach. Despite the overall resemblance, the derived model reveals some key differences between BRD8 and BRD9 in amino acid identities proximal to the binding pocket (Figure 3A). Among these differences, we noticed in particular that BRD9’s Tyr “gatekeeper” residue (**Y222**), which separates the KAc pocket from the ZA shelf (otherwise known as the “WPF” shelf within the BET family), is replaced with a more sterically permissive valine within the BRD8 structure (**V793**) (Figure 3B, 3C). Likewise, the nearby BRD9 shelf Phe residue (**F160**), is also replaced with a more compact valine (**V731**). We considered that these amino acid substitutions might accommodate substituents projecting toward this region from BI-9564’s biaryl ortho positions (Figure 3C, Arrow), which would otherwise introduce a steric clash with Y222/F160 within the BRD9 domain. Therefore, in pursuit of BRD8 selectivity, we designed analogs that featured a 2,6-disubstitution pattern order to enforce projection into this region. Encouragingly, compound **<u>2</u>**, featuring a fused phenyl ring and ortho methoxy, and compound **<u>3</u>**, featuring a symmetrical 2,6-dimethoxy substitution, both showed dramatically reduced activity against BRD9 (>10µM IC50), while retaining activity against BRD8 (Table 2). In fact, compound **<u>3</u>** showed an 11-fold increase in BRD8 potency (IC50 = 85 nM) versus parent BI-9564 (Table 2).

**Table 2.** Gatekeeper directed steric sussitutions.

|  | Cmpd 2 |  |
| --- | --- | --- |
| | BRD8(1) ($\mu\text{M}$) | 0.40 |
| | BRD9 ($\mu\text{M}$) | >10 $\mu\text{M}$ |
| | BRD4(1) ($\mu\text{M}$) | >10 $\mu\text{M}$ |
|  | Cmpd 3 |  |
| | BRD8(1) ($\mu\text{M}$) | 0.085 |
| | BRD9 ($\mu\text{M}$) | >10 $\mu\text{M}$ |
| | BRD4(1) ($\mu\text{M}$) | >10 $\mu\text{M}$ |

**Fig 3.**
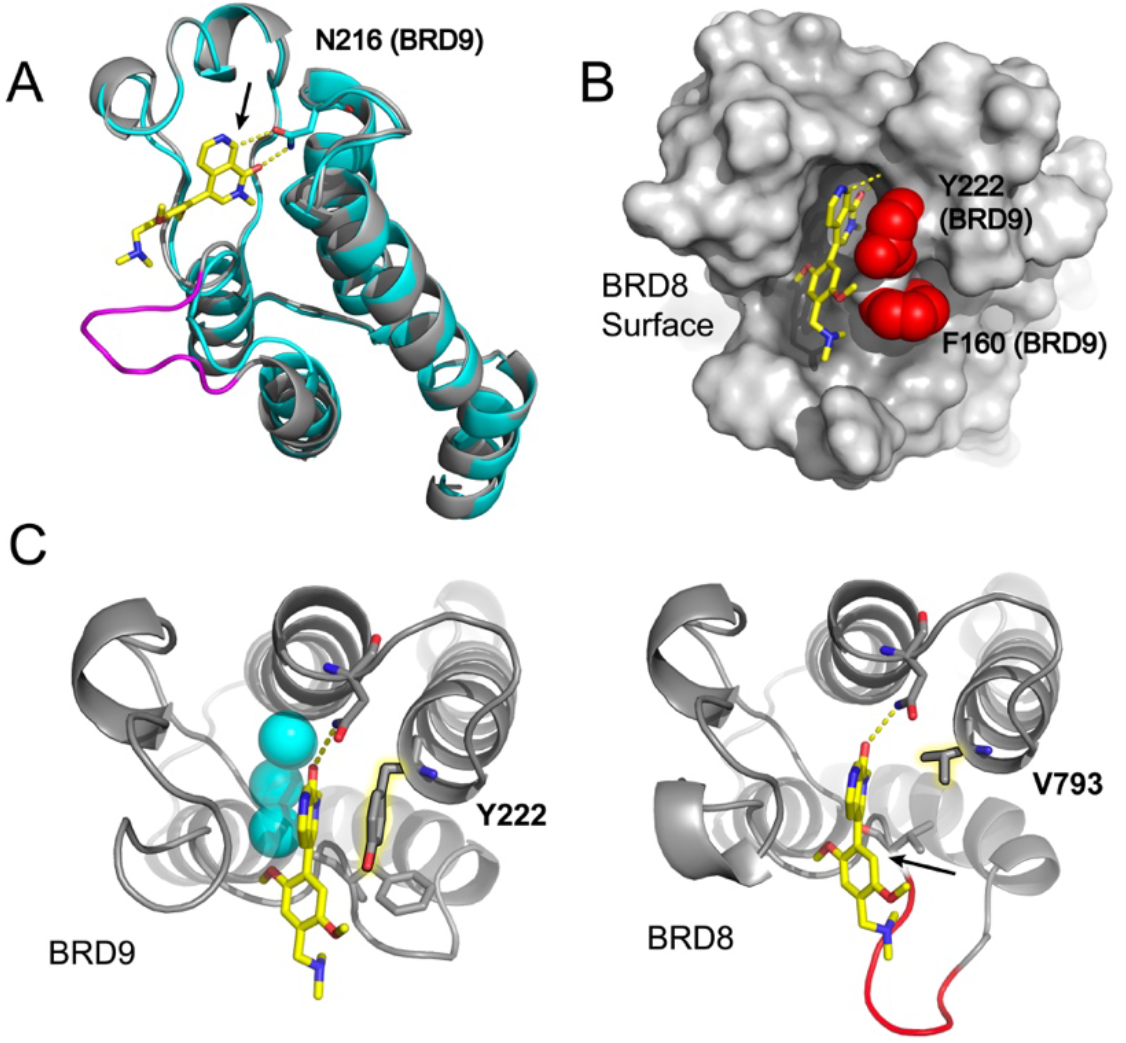
Homology model guided design toward potent and selective BRD8(1) bromodomain binders. (A) Overlay of BRD8(1) homology model (cyan cartoon) with published BRD9 bromodomain-BI-9564 co-crystal structure (gray cartoon), highlighting BRD8 polar insertion region (magenta) and N216 H-bonding within BRD9 (dashed). Arrow demarks position of naphthyridone polar contact. (B) Overlay as in A, showing BRD8(1) surface with docking of BI-9564. Adjacent BRD9 gatekeeper Y222 and Shelf F106 residues are highlighted (red spheres). (C) Cartoon display of BRD9-BI-9564 cocrystal (left) and BRD8(1)-BI-9564 docked homology model (right) with highlighted gatekeeper residues (glowing sticks), and ASN H-bonding (dashed). Arrow demarks shelf projecting substituent position.

In a parallel strategy toward improving potency, we focused our attention on an expected polar contact from naphthyridone 8 position aryl hydrogen to the KAc anchoring Asn residue (BRD8 **N791**) (Figure 3A, Arrow). Here we envisioned that alternative electron withdrawing groups within the head group’s embedded pyridine ring might serve to tune this interaction to optimize BRD8 affinity. We observed that substitution of the embedded pyridine for an electron deficient pyrimidine resulted in reduced activity (**<u>4</u>**), as did replacement of the pyridyl nitrogen with highly withdrawing nitro and trifluoromethyl ring substituents (**<u>5,6</u>**). We found, however, that aminocarbonyl substituted compound **<u>7</u>** was approximately equipotent to BI-9564, while cyano and fluoro substituted compounds (**<u>8, 9</u>**) showed significantly improved BRD8 activity (Table 3). This SAR affirmed our expectations for this position as an interaction hotspot, although importantly, given its position within the rim of the KAc pocket, we suspect these modifications contribute to both the intended electronic effects on aryl polar contact donation, as well as to direct substituent contributions to binding. Together, our initial SAR studies identified hotspots for tuning BRD8 potency and selectivity, established the suitability of the BI-9564 scaffold for development into high-quality probes, and provided an encouraging validation of our BRD8 homology model.

**Table 3.**
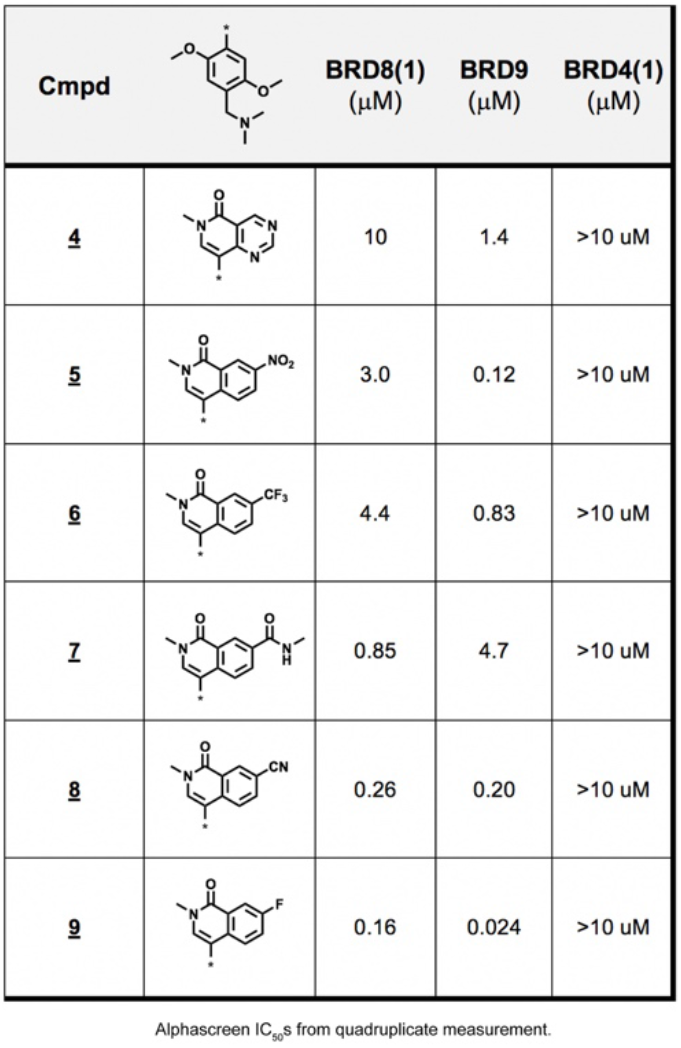
ASN H-bonding directed electron withdrawing group modifications.

| Cmpd | | BRD8(1) ($\mu\text{M}$) | BRD9 ($\mu\text{M}$) | BRD4(1) ($\mu\text{M}$) |
| --- | --- | --- | --- | --- |
| **4** | | 10 | 1.4 | >10 $\mu\text{M}$ |
| **5** | | 3.0 | 0.12 | >10 $\mu\text{M}$ |
| **6** | | 4.4 | 0.83 | >10 $\mu\text{M}$ |
| **7** | | 0.85 | 4.7 | >10 $\mu\text{M}$ |
| **8** | | 0.26 | 0.20 | >10 $\mu\text{M}$ |
| **9** | | 0.16 | 0.024 | >10 $\mu\text{M}$ |

Alphascreen $\text{IC}_{50}$ s from quadruplicate measurement.

Having identified avenues towards more potent and selective BRD8 inhibitors, we next sought to convert our initial SAR insights toward tool development for an in-cell BRD8 engagement assay using the previously described BRET tracer displacement format.^25^ Our previous attempts in this regard had met limited success due the poor potency of fluorescent tracers based on BI-9564. We therefore incorporated the bromodomain warhead of compound **<u>3</u>**, the most potent BRD8 binder of our initial series, to prepare a NanoBRET tracer conjugate (Figure 4A). Pairing this tracer with BRD8(1) expressed as a fusion with nanoluciferase in HEK293T cells, we were able to establish a robust, dose-responsive BRET assay, which alongside similarly constructed BRD9 and BRD4 assays, provided a bromodomain target engagement panel in live cells. (Figure 4A, 4B) (Table 4). The in-cell BRET activity of literature bromodomain probes was consistent with the in-vitro AlphaScreen profiles, corroborating relative potency rankings of this probe set against BRD9 and BRD4, as well as confirming BI-9564 to be the strongest binder of BRD8 (Table 4).

With a suite of tools for evaluation of BRD8 bromodomain inhibitors in hand, we aimed to apply insights from our prior chemical series in hopes of arriving at high quality probe compounds. Incorporation of the top performing fluoro-isoquinolone head group of compound **<u>9</u>**, alongside the gatekeeper-directed 2,6-dimethoxy configuration of compound **<u>3</u>**, gave **DN01** and paired chemoproteomics tool compound **DN02** (Figure 5A). Both **DN01** and **DN02** showed low nM AlphaScreen IC50s (12 nM & 48 nM respectively), with high selectivity over BRD9 and BRD4 in biochemical AlphaScreen assays, and in-cell engagement within BRET displacement assays (Figure 5A, 5B). The DiscoveRx phage-display bromoKdELECT assay further confirmed the potency of **DN02** against BRD8(1) (32 nM), and interestingly, demonstrated this activity to be selective over BRD8’s second bromodomain (BRD8(2) >1000 nM) (Supplemental Figure 1). Docking **DN01** into the BRD8(1) homology model illustrated a binding mode analogous to that observed for BI-9564 to BRD9, and indicated the 2,6-dimethoxy substitution to project towards the permissive gatekeeper valine in BRD8 as desired (Figure 5C). A comparative docking of BI-9564 into BRD8’s first and second bromodomains revealed substitution of two BRD8(1) residues that provide critical hydrophobic contacts to the **DN01** phenyl ring (**V731** to **P1127** and **I740** to **Q1136**), providing insight into the reduced potency against BRD8(2) (Supplemental Figure 2). We expect that substitution of BRD8(1)’s Ile740 for hydrophilic Glu1136 within BRD8(2) particularly accounts for the reduced binding affinity.

**Table 4.**
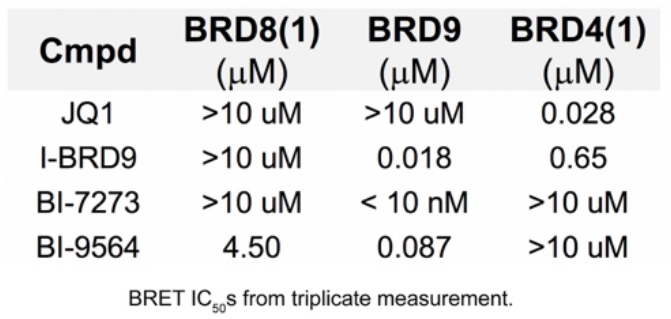
Characterization of Literature Compounds by In-Cell BRET Assay.

**Fig 4.**
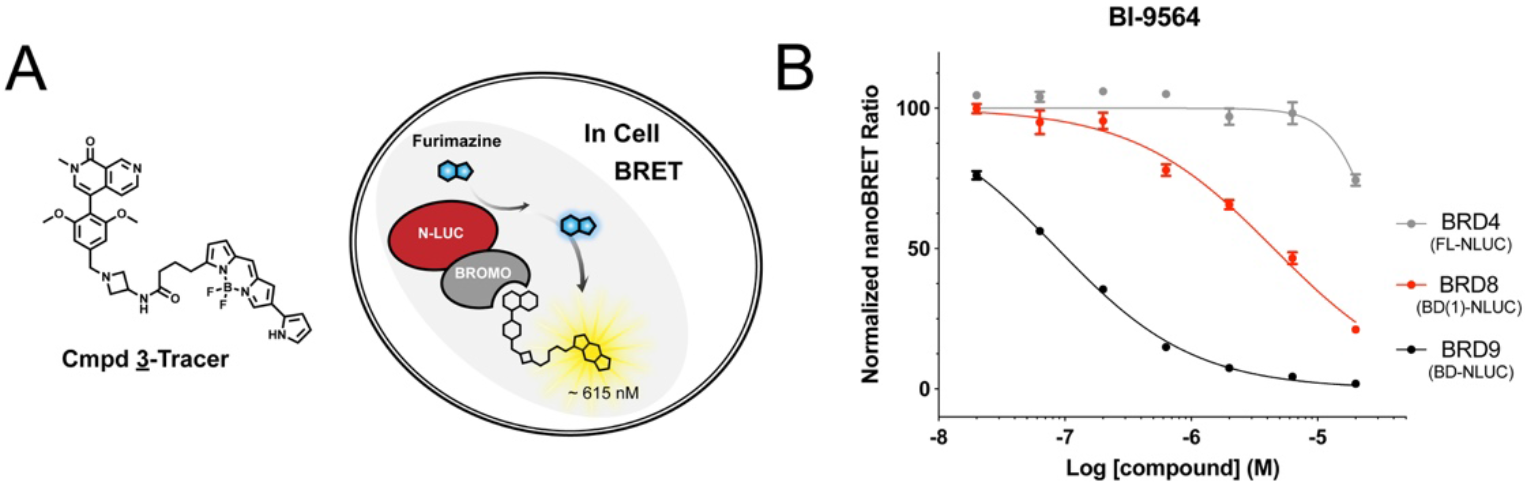
Implementation of a Cellular BRET assay to confirm compound binding in live cells. (A) Structure of BRD8 BRET tracer (Compound <u>3</u>-Tracer) (left) and assay schematic (right). (B) Representative In-Cell BRET competition curves for literature compound BI-9564.

**Fig 5.**
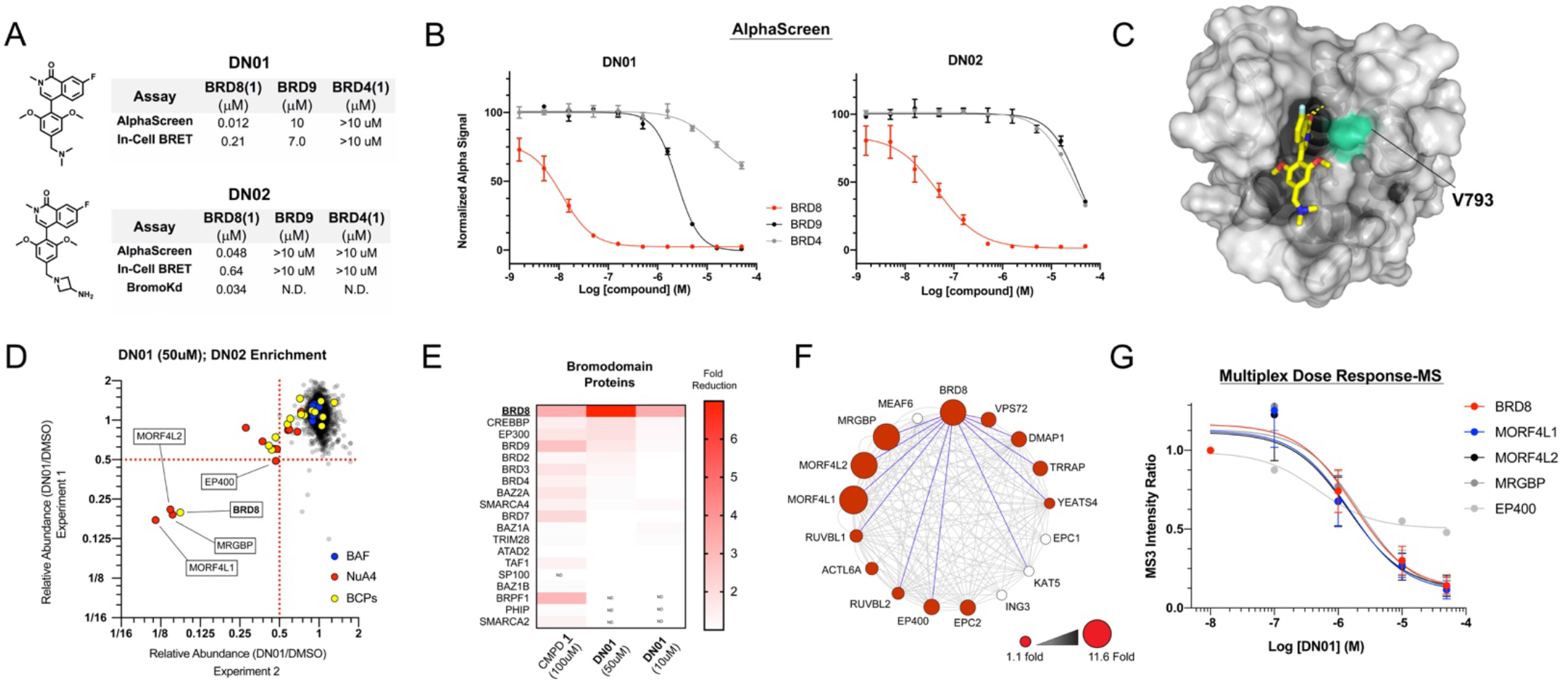
Characterization of selective BRD8 bromodomain probes DN01 and DN02. (A) Structures and binding data for DN01 and DN02 and embedded tabulated data for indicated assays. (B) Representative AlphaScreen competition curves for DN01 and DN02. (C) Docking of DN01 within BRD8(1) homology model with permissive gatekeeper V793 highlighted (cyan spheres). (D) Scatterplot view of duplicate DN02 enrichment chemoproteomics with DN01 competition (50µM). NuA4 / BAF complexes and bromodomain-containing protein (BCP) targets are highlighted. (E) Heatmap of fold competition across all identified BCP targets in DN01 and Compound <u>1</u> datasets (ranked by 50µM DN01 data). (F) Co-competition of NuA4 complex members (CORUM database) within 50µM DN01 chemoproteomics data. Fold competition is indicated by circle radius for identified members (red circles), network connectivity within the BioPlex database is shown with connecting lines, and BRD8 bait interactions are highlighted (blue lines). (G) Competitive intensity ratio curves for DN02 enrichment chemoproteomics multiplexed across a concentration range of DN01. Curves are shown for proteins identified with >50% competition at 50µM.

To assess the proteome-wide binding profile of our advanced BRD8(1) bromodomain ligands, we returned to our chemoproteomic profiling approach, performing enrichment with a **DN02** functionalized matrix in the presence of free **DN01** as a competitor. Here, we applied 10-channel TMT multiplexing across increasing concentrations of **DN01** in order to resolve dose response relationships. Consistent with in-vitro binding results, 50μM **DN01** selectively competed BRD8 and an assembly of additional NuA4 complex members, while retaining marked selectivity over activity on BRD9 and associated BAF complex members (Figure 5D). Indeed, five members of the NuA4 complex were the only proteins competed by at least twofold in both technical replicates. Accordingly, **DN01** showed selectivity within the bromodomain family, with no other bromodomain-containing proteins showing complete dose-responsive curves up to the top dose (50μM). Although we note initial apparent dose-responsive competition for CBP, P300 and BRD9, this activity was insufficient to derive curve fits, indicating low affinity interactions (Figure 5E). These observations were supported by a DiscoveRx BromoMax competition panel, with **DN02** showing minimal off-target engagement across the bromodomain family, with modest activity against CBP/P300 (Supplemental Table 1) (BRD8 currently available in dose response format only: Kd=32 nM).

Within the NuA4 complex, our **DN01**/**DN02** chemoproteomics dataset identified a total of 13 of the 17 complex subunits, as collated by the CORUM database (Figure 5F). Of these identified members, subunits previously annotated as BRD8 interactors within the BioPlex affinity purification interactome dataset showed excellent representation, with 10 out of 11 previously reported interactors detected. Among these previously identified BRD8 interactors were the five top **DN01** competed NuA4 members: BRD8, MORFL1/2, MRGBP, and EP400 (>2 fold competition, two replicates). These proteins showed robust and coordinated competition profiles across multiplexed concentrations of DN01, reflecting engagement with the intact NuA4 complex via binding to BRD8 (Figure 5G). Collectively, these data nominate **DN01**/**DN02** as a qualified chemical probe for the selective engagement of BRD8 and enrichment of native NuA4 complexes.

## Discussion

In this work, we investigate the proteome-wide binding profile of a BAF-engaging BRD9 inhibitor scaffold using chemical affinity proteomics. Unexpectedly, we discover engagement of the NuA4 acetyltransferase complex by a derivative of this well-studied series (Compound **1**), which we link to a previously unreported binding interaction with the bromodomain-containing subunit BRD8. Despite compelling rationale for study, BRD8 has remained inaccessible by reported chemical discovery campaigns. Having eluded crystallographic characterization, and lacking reported binding assays, evidence of BRD8 druggability has been thus far limited to promiscuous covalent warhead studies.^24^ Leveraging the uncovered compound **1** -BRD8(1) interaction, we develop in-vitro, cellular, and chemoproteomics assays for BRD8(1) interaction. Finally, implementing these tools, we apply model-guided design to elaborate first-in-class selective, ligand efficient, and cellularly active compounds targeting BRD8(1). Importantly, although these findings were empowered by unexpected BRD8 off-target activity of literature probes, we note that there currently is no evidence that this relatively weak activity meaningfully contributes to the reported phenotypes of these potent BRD9 inhibitors. Indeed, this interaction did not previously support targeted degradation of BRD8 when BRD9 ligands were adapted as CRBN-recruiting degraders.^6^

The current findings promote the BRD8 bromodomain as an accessible target for discovery chemistry and establish a first chemical foothold within the NuA4 acetyltransferase complex. The predictive success of BRD9-templated homology modeling to our experimental SAR supports a conserved fold for BRD8, despite an atypical four residue polar insertion proximal to the KAc in the homologous ZA loop region. In particular, we exploit the predicted positioning of a sterically permissive Val gatekeeper residue as a primary means to discriminate activity against BRD9 and other parental ligand targets, enabling the development of BRD8 selective probes **DN01** and **DN02**. Notably, profiling of **DN02** against both domains of BRD8 further reveals a dramatic preferential binding to BD1. Considering that studies have demonstrated biologically distinct outcomes for mono-specific inhibitors of other multi-bromodomain targets, this feature may offer an advantage in biological specificity.^26-27^

The probes resulting from this work provide new opportunities for the study of the role of BRD8’s bromodomain within the NuA4 complex, as well as ligands for affinity enrichment and targeted degradation approaches. In this regard, an accumulating body of work has implicated a tumor-supportive role for BRD8 in colorectal carcinoma,^17, 19-20^ and more recently in hepatocellular carcinoma,^21^ characterized by its frequent overexpression, correlation with poor prognosis, and contribution to traditional chemotherapy resistance.^20^ These studies link this biology to BRD8’s influence in DNA repair pathways and associated signaling, consistent with the longstanding appreciation of a role for NuA4 and the TIP60 acetyltransferase activity in these pathways.^18, 28^ The tools developed herein will enable assessment of bromodomain inhibitors’ ability to recapitulate antiproliferative effects of genetic knockdown of BRD8 in these contexts,^17, 20-21^ or if further adaptation as targeted protein degradation probes will be required. Finally, the findings of this study further highlight the power of chemoproteomics workflows to provide both unbiased interrogation of target space and comparative assessment of complex engagement. The broad assembly of chromatin regulatory proteins into multisubunit complexes suggests these targets to be a fruitful class for a continued application of this strategy.

## Acknowledgements

The authors thank Catherine Dubreuil for her efforts directing and connecting the Harvard Therapeutics industry internship, as well as the HITS leadership for their forward thinking program. We also thank Michael Salcius for his support with protein expression, Rishi Jain and John Tallarico for helpful discussions and support of open publication, and Jay Bradner for his promotion of open science and collaborative opportunities at NIBR.

## Supplemental

**Table S1.**
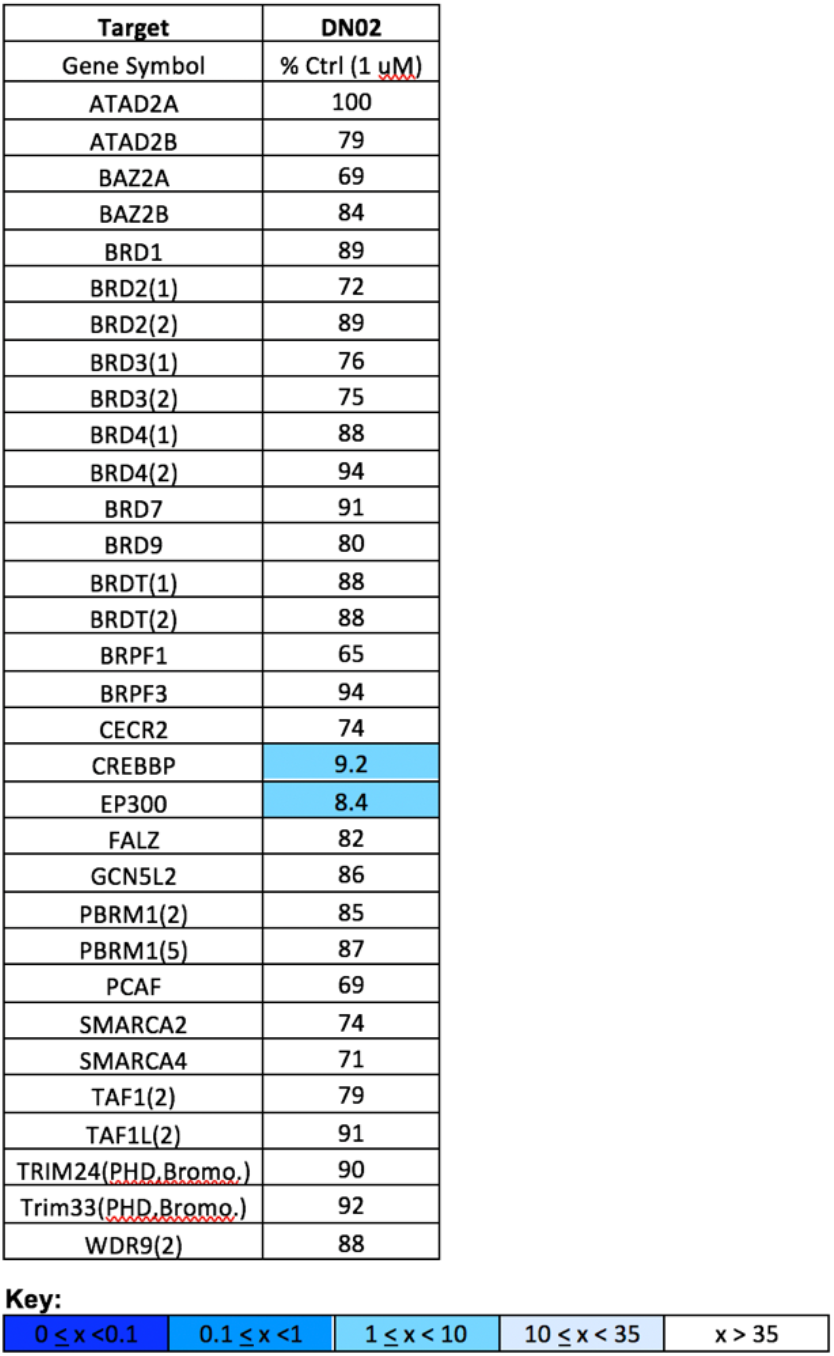
Single point screening of **DN02** off targets at 1µMusing Bromoscan (DiscoveRx).

| Target | DN02 |
| --- | --- |
| Gene Symbol | % Ctrl (1 μM) |
| ATAD2A | 100 |
| ATAD2B | 79 |
| BAZ2A | 69 |
| BAZ2B | 84 |
| BRD1 | 89 |
| BRD2(1) | 72 |
| BRD2(2) | 89 |
| BRD3(1) | 76 |
| BRD3(2) | 75 |
| BRD4(1) | 88 |
| BRD4(2) | 94 |
| BRD7 | 91 |
| BRD9 | 80 |
| BRDT(1) | 88 |
| BRDT(2) | 88 |
| BRPF1 | 65 |
| BRPF3 | 94 |
| CECR2 | 74 |
| CREBBP | 9.2 |
| EP300 | 8.4 |
| FALZ | 82 |
| GCN5L2 | 86 |
| PBRM1(2) | 85 |
| PBRM1(5) | 87 |
| PCAF | 69 |
| SMARCA2 | 74 |
| SMARCA4 | 71 |
| TAF1(2) | 79 |
| TAF1L(2) | 91 |
| TRIM24(PHD,Bromo.) | 90 |
| Trim33(PHD,Bromo.) | 92 |
| WDR9(2) | 88 |

Key:
|  |  |  |  |  |
| --- | --- | --- | --- | --- |
| $0 \leq x < 0.1$ | $0.1 \leq x < 1$ | $1 \leq x < 10$ | $10 \leq x < 35$ | $x > 35$ |

**Figure S1.**
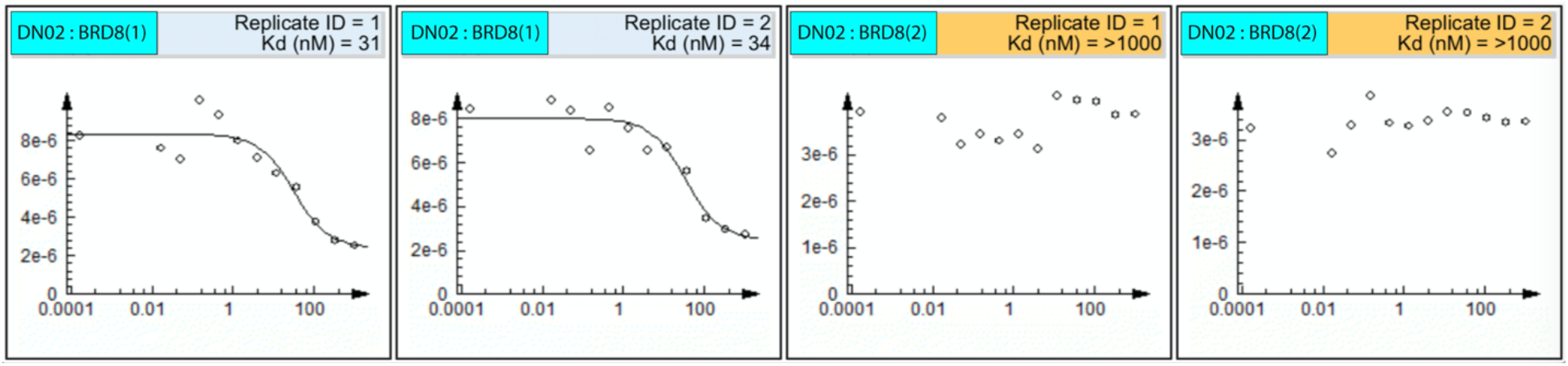
Dose response BromoKD curves for DN02 (DiscoveRx).

**Figure S2.**
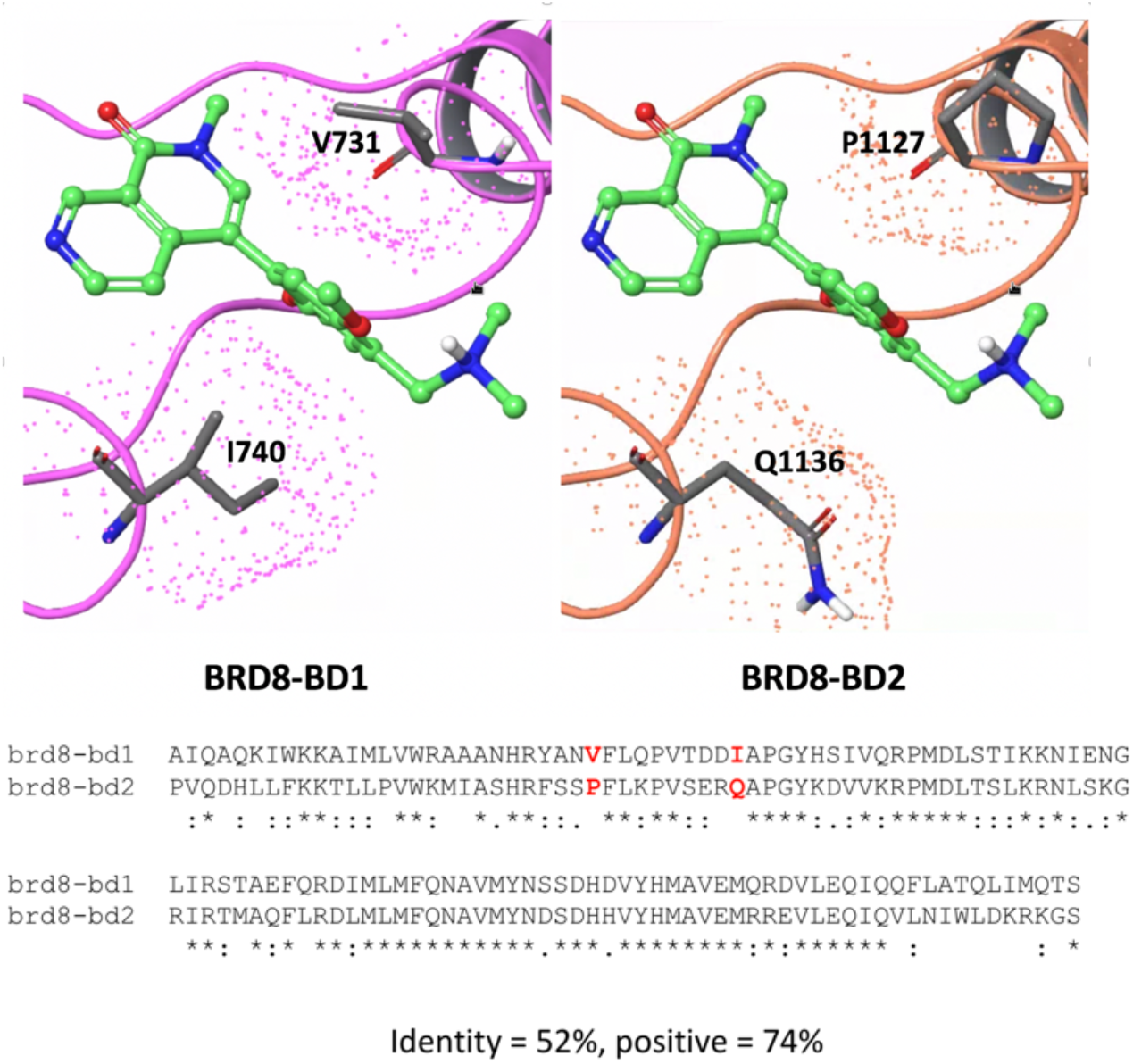
(Top) Comparison of BRD8(1) and BRD8(2) homology model structures with docked ligand (BI-9564). Key proximal residue differences are labeled. (Bottom) BRD8(1) and BRD8(2) sequence alignment.

